# Gene regulation of infection-associated L-tartrate metabolism in *Salmonella enterica* serovar Typhimurium

**DOI:** 10.1101/2024.02.05.578992

**Authors:** Vivian K. Rojas, Maria G. Winter, Angel G. Jimenez, Natasha W. Tanner, Stacey L. Crockett, Luisella Spiga, David R. Hendrixson, Sebastian E. Winter

## Abstract

Enteric pathogens such as *Salmonella enterica* serovar Typhimurium experience spatial and temporal changes to the metabolic landscape throughout infection. Host reactive oxygen and nitrogen species non-enzymatically convert monosaccharides to alpha hydroxy acids, including L-tartrate. *Salmonella* utilizes L-tartrate early during infection to support fumarate respiration, while L-tartrate utilization ceases at later time points due to the increased availability of exogenous electron acceptors such as tetrathionate, nitrate, and oxygen. It remains unknown how *Salmonella* regulates its gene expression to metabolically adapt to changing nutritional environments. Here, we investigated how the transcriptional regulation for L-tartrate metabolism in *Salmonella* is influenced by infection-relevant cues. L-tartrate induces the transcription of *ttdBAU*, genes involved in L-tartrate utilization. L-tartrate metabolism is negatively regulated by two previously uncharacterized transcriptional regulators TtdV (STM3357) and TtdW (STM3358), and both TtdV and TtdW are required for sensing of L-tartrate. The electron acceptors nitrate, tetrathionate, and oxygen repress *ttdBAU* transcription via the two-component system ArcAB. Furthermore, regulation of L-tartrate metabolism is required for optimal fitness in a mouse model of *Salmonella*-induced colitis. TtdV, TtdW, and ArcAB allow for the integration of two cues, substrate availability and availability of exogenous electron acceptors, to control L-tartrate metabolism. Our findings provide novel insights into how *Salmonella* prioritizes utilization of different electron acceptors for respiration as it experiences transitional nutrient availability throughout infection.

**IMPORTANCE:** Bacterial pathogens must adapt their gene expression profiles to cope with diverse environments encountered during infection. This coordinated process is carried out by the integration of cues that the pathogen senses to fine-tune gene expression in a spatiotemporal manner. Many studies have elucidated the regulatory mechanisms on how *Salmonella* sense metabolites in the gut to activate or repress its virulence program, however our understanding of how *Salmonella* coordinates its gene expression to maximize the utilization of carbon and energy sources found in transitional nutrient niches is not well understood. In this study, we discovered how *Salmonella* integrates two infection-relevant cues, substrate availability and exogenous electron acceptors, to control L-tartrate metabolism. From our experiments, we propose a model for how L-tartrate metabolism is regulated in response to different metabolic cues in addition to characterizing two previously unknown transcriptional regulators. This study expands our understanding of how microbes combine metabolic cues to enhance fitness during infection.

## INTRODUCTION

Enteric pathogens coordinate gene expression in a spatial manner to ensure production of virulence factors in the correct anatomical location within the host. Spatial sensing is achieved by a plethora of cues, many of which are either host- or microbiota-derived metabolites. For example, the Enterohaemorrhagic *E. coli* and *Citrobacter rodentium* sense microbiota-derived galacturonic acid through ExuR, which in turn activates the expression of the type three secretion system (T3SS) genes found on the LEE pathogenicity island (1). Fucose, another metabolite liberated by the microbiota, is sensed by Enterohaemorrhagic *E. coli* to activate the two-component system FusKR which in turn controls the expression of virulence and metabolic genes (2). Host-derived oxygen and microbiota-derived formate regulate the virulence program of *Shigella flexneri*, an intracellular pathogen that invades the colonic epithelium (3). The invasion-associated T3SS (T3SS-1) in *Salmonella enterica* serovar Typhimurium (*S*. Tm) is regulated by a large number of metabolic cues, including formate, butyrate, acetate and long chain fatty acids (4–8). This complex regulatory network ensures expression of T3SS-1 in the distal small intestine, the primary site of invasion (9). As such, the location-specific sensing of metabolites is critical for the expression of virulence factors and tissue-specific induction of disease.

The local inflammatory response induced by invasive enteric pathogens, such as *S*. Tm, suppresses host-associated microbial communities in the gut while facilitating expansion of a pathogen population in the gut lumen (10), which in turn enhances transmission success by the fecal-oral route (11). The expansion of the *S*. Tm population is driven by changes in the metabolic landscape as inflammation develops (12). In the initial stage of infection, *Salmonella* utilizes simple sugars, amino acids, and molecular hydrogen (13–15). Upon tissue invasion, *Salmonella* induces localized inflammatory responses, and transmigrating phagocytes undergo an oxidative burst to release reactive oxygen species and nitric oxide radials. These phagocyte-derived compounds react to form nitrate or interact with endogenous molecules such as thiosulfate to form tetrathionate (16, 17). Furthermore, a change in epithelial cell metabolism results in the diffusion of molecular oxygen into the gut lumen (18). A lumenal population of *Salmonella* performs respiration using nitrate, tetrathionate, and oxygen to promote an oxidative central metabolism at later stages of infection (19). Using respiration, *Salmonella* utilizes a variety of poorly fermentable metabolites, thus bypassing nutritional competition with anaerobic members of the microbiota (19–23). While several metabolic pathways used by *S*. Tm for gut colonization have been described, our understanding of how *S*. Tm responds to temporal changes in metabolite availability is limited.

We have recently shown that at early time points, prior to the overt transmigration of neutrophils, release of reactive oxygen species and nitric oxide radicals leads to a non-enzymatic decarboxylation of monosaccharides (24). This reaction generates alpha hydroxy carboxylic acids, including the tartrate isomers L- and D-tartrate. *Salmonella* utilizes L- and D-tartrate in a stereospecific manner (24). After uptake of L-tartrate by the TtdU transporter, the dehydratase TtdAB converts L-tartrate to oxaloacetate. Part of the oxaloacetate pool enters the reductive branch of the TCA cycle to support fumarate respiration, while a portion of oxaloacetate is converted to pyruvate via the oxaloacetate decarboxylase sodium pump. Curiously, utilization of L-tartrate only occurs at earlier time points during infection, and apparently, L-tartrate utilization ceases once overt inflammation has developed (24). Experiencing transient nutrient niches requires an adaptation to the changing environments, and these adaptations are influenced by gene expression. In this study, we investigate the gene regulation for *Salmonella* L-tartrate metabolism as it relates to infection-relevant cues.

## RESULTS

### Substrate induction of the L-tartrate utilization gene cluster

The *ttdBAUVW* gene cluster in *S*. Tm encodes for the two subunits of the L-tartrate dehydratase, TtdA and TtdB, the transporter TtdU, and two GntR-type regulators, STM3357 (renamed TtdV) and STM3358 (renamed TtdW) (**Fig. 1A**). Since genes involved in catabolism are commonly regulated by substrate availability, we initially investigated gene expression of the *ttd* gene cluster in response to L-tartrate. To this end, we created a transcriptional fusion of the *ttdA* gene and *lacZ* (encoding β-galactosidase). The transcriptional fusion did not interfere with L-tartrate utilization (**Fig. S1A**). We first quantified β-galactosidase activity in response to increasing concentrations of L-tartrate (**Fig. 1B**). The half maximal effective concentration (EC_50_) for L-tartrate was approximately 0.1 mM. Neither D-nor *meso*-tartrate elicited β-galactosidase activity, suggesting that transcriptional induction by L-tartrate is stereospecific.

**Figure 1:**
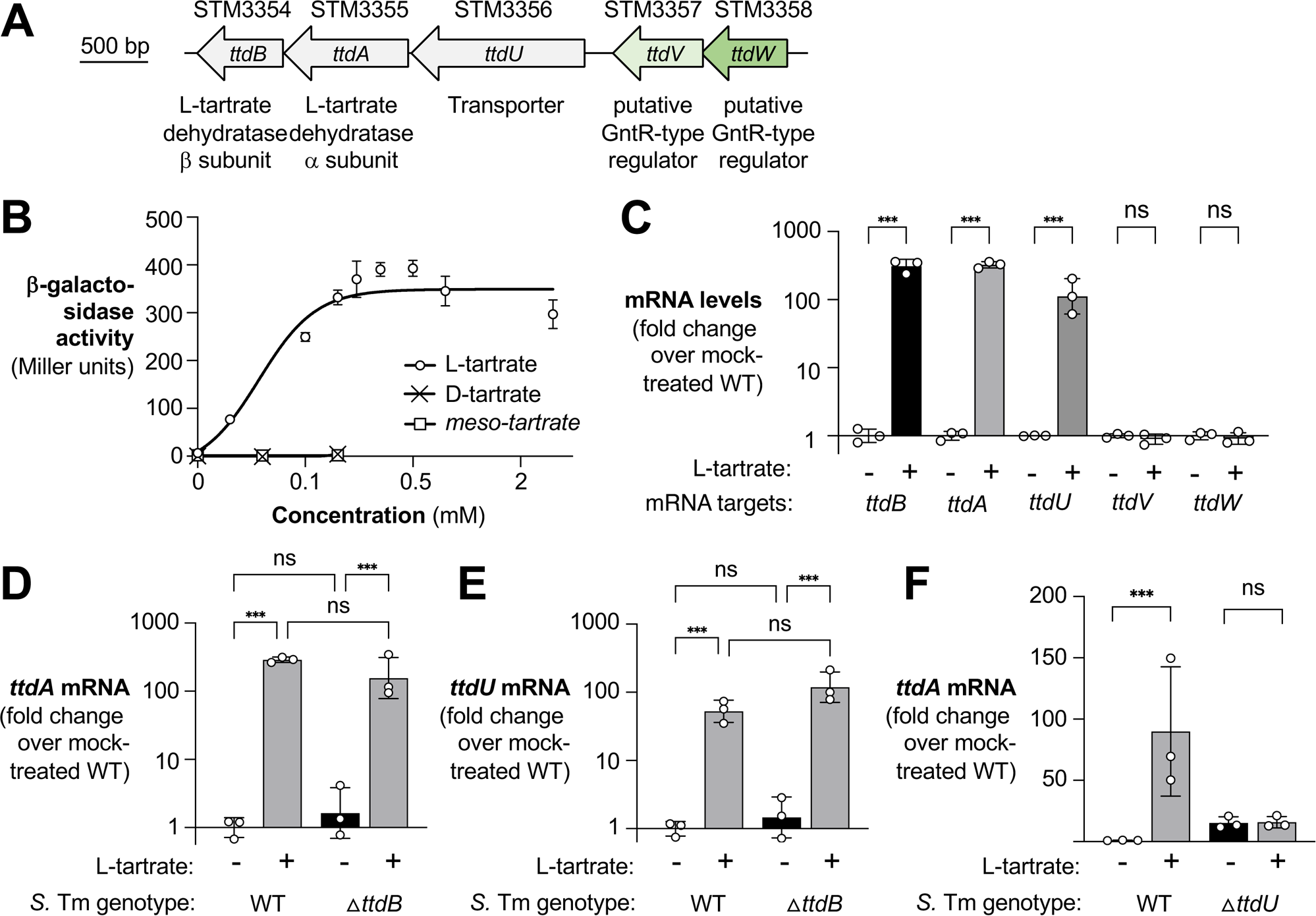
Substrate induction of the L-tartrate utilization gene cluster. **(A)** Schematic representation of the *ttdBAU-ttdVW* gene cluster in *S.* Tm LT2. The gene locus number and (putative) function of the gene product is listed. STM3357 and STM3358 were renamed *ttdV* and *ttdW* consistent with their proposed function of regulating L-tartrate metabolism in *S*. Tm. **(B)** The transcriptional *ttdA-lacZ* fusion strain (*ttdA*::pVR7) was cultured in media containing M9 salts and casamino acids (M9C) supplemented with various concentrations of L-, D- or *meso-tartrate* and grown anaerobically. After 5 hours, β-galactosidase activity was assessed. **(C)** The *S*. Tm wild-type strain was grown anaerobically in M9C media supplemented without or with 0.1 mM L-tartrate for 3 hours. Total RNA was extracted and mRNA levels quantified by RT-qPCR, mRNA levels were normalized to the house keep gene *gmk*. (D - E) mRNA levels of *ttdA* **(D)** and *ttdU* **(U)** were determined in the wild-type strain and the *ttdB* mutant, cultured in the absence (black bars) and presence of L-tartrate (gray bars). **(F)** mRNA levels of *ttdA* were determined in the wild-type strain and the *ttdU* mutant. Bacteria were cultured in the absence (black bars) and presence of L-tartrate (gray bars). Bars represent the geometric mean with geometric standard deviation. Each dot represents one biological replicate. WT; wild-type *** *P* < 0.001; ns, not significant (ANOVA)

Next, we analyzed the transcription of each gene in the *ttd* gene cluster in the presence or absence of L-tartrate. Media, supplemented with 0.1 mM L-tartrate (EC_50_) as indicated, was inoculated with the *S*. Tm wild-type strain. After 3 h of growth under anaerobic conditions, we extracted RNA and quantified mRNA levels by RT-qPCR (**Fig. 1C**). mRNA levels of the transporter (*ttdU*) and the two subunits of the L-tartrate dehydratase (*ttdA* and *ttdB*) were induced by more than 100-fold in response to L-tartrate. In contrast, *ttdV* and *ttdW* mRNA levels were unchanged (**Fig. 1C**), even at higher, likely physiological irrelevant (24) concentrations of L-tartrate (**Fig. S1B**).

We considered that, rather than L-tartrate, a metabolite derived from L-tartrate degradation might be a cue for induction of *ttdBAU* transcription. To test this idea, we generated a mutant lacking L-tartrate dehydratase activity (*ttdB* mutant) and determined *ttdA* and *ttdU* mRNA levels (**Fig. 1D and E**). L-tartrate stimulated *ttdA* and *ttdU* transcription even in the absence of L-tartrate dehydratase activity. Furthermore, induction by L-tartrate was absent in a *ttdU* mutant (**Fig. 1F**), consistent with the idea that L-tartrate must enter the cell to influence gene regulation. Collectively, these experiments suggest that L-tartrate is an inducer of *ttdBAU* transcription.

### TtdV and TtdW are transcriptional repressors for the L-tartrate utilization genes

TtdV and TtdW are members of the GntR family of bacterial transcriptional regulators. These types of transcriptional regulators have a conserved DNA-binding motif at the N-terminus, while the C-terminus serves as a ligand binding or oligomerization domain (25). Transcriptional regulators in the GntR family can be either transcriptional activators or repressors (26). To determine the function of TtdV and TtdW, we created clean deletion of the *ttdV*, *ttdW*, and *ttdVW* genes. Cultivation in Jordan’s Tartrate Agar, a growth medium designed to detect fermentation of tartrate, indicated robust tartrate utilization by the *ttdV*, *ttdW*, and *ttdVW* mutants (**Fig. S2**). Transcription of *ttdA* was significantly elevated in these mutants compared to the wild-type strain and these strains were non-responsive to L-tartrate (**Fig. 2A**), suggesting that TtdV and TtdW are negative regulators of *ttdBAU* gene regulation. A *ttdVW* double mutant exhibited a phenotype similar to the single mutant. Genetic complementation, either by introducing the *ttdV* or *ttdW* coding sequence and its native promoter into the neutral *phoN* locus or by expressing recombinant histidine-tagged fusion proteins, restored repression of the L-tartrate utilization genes under non-inducing conditions (**Fig. 2A and S2B**). Furthermore, we generated mutants in which we replaced the start codon of *ttdV* (1GTG>GTT) and *ttdW* (1GTG>TAG). One would predict that with this mutagenesis strategy, any potential *cis*-regulatory elements concealed in the *ttdV* or *ttdW* coding sequence should be largely retained. Akin to the single gene deletions, these mutants exhibited increased *ttdA* transcription in the absence of L-tartrate (**Fig. 2B and C**). Collectively, these experiments indicate that TtdV and TtdW are repressors of *ttdBAU,* and L-tartrate relieves repression by TtdV and TtdW.

**Figure 2:**
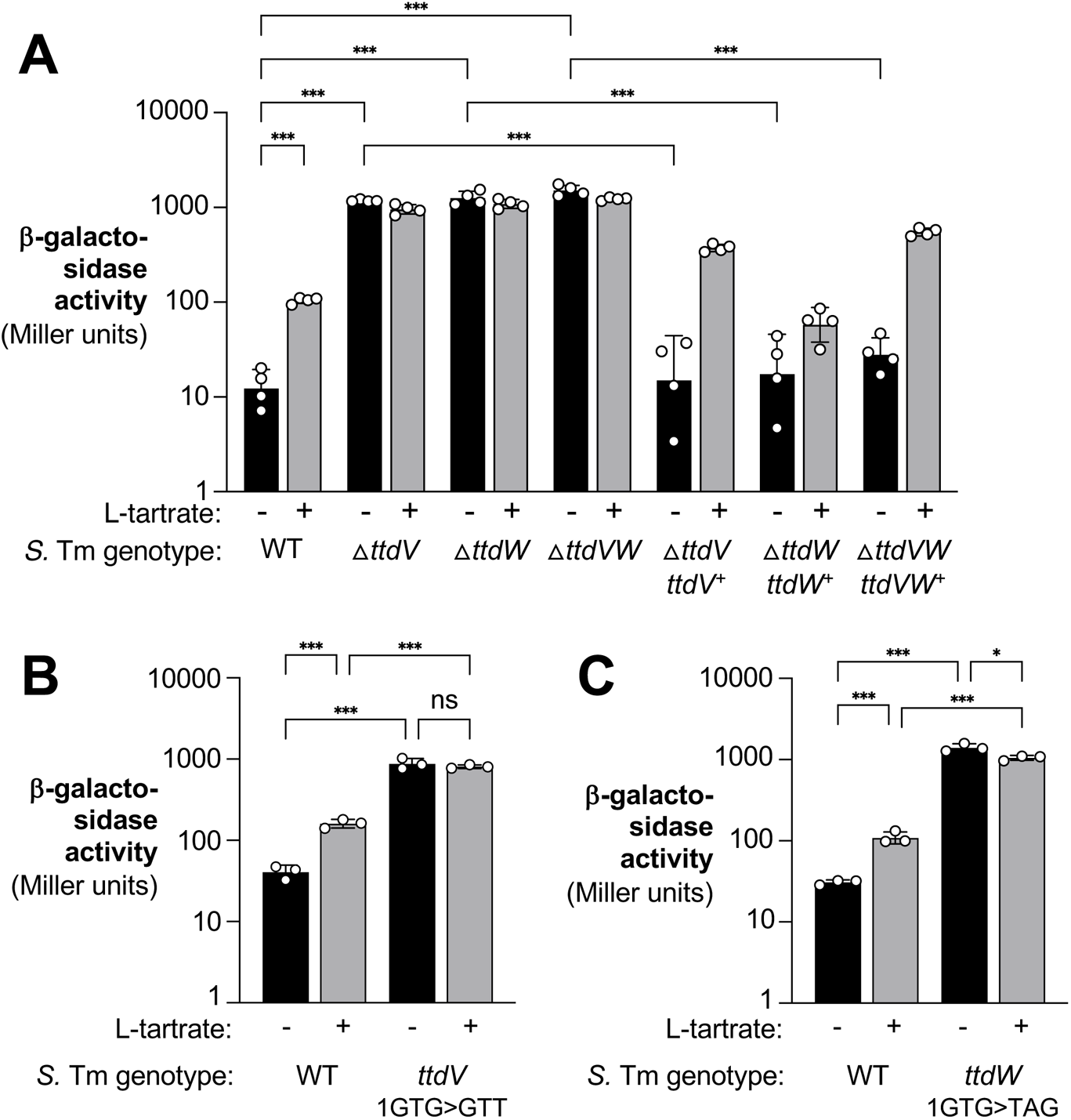
TtdV and TtdW are transcriptional repressors for the L-tartrate utilization genes. **(A)** The indicated *S.* Tm *ttdA-lacZ* transcriptional fusion strains were grown anaerobically without (black bars) or with L-tartrate (0.1 mM; gray bars). After 3 hours, β-galactosidase activity was determined. The pertinent genotype is listed. For genetic complementation, genes of interest and their native promoters were inserted into the neutral *phoN* gene. **(B and C)** The start codon of the *ttdV* gene **(B)** and the *ttdW* gene **(C)** were altered as indicated. Bacteria carrying a *ttdA-lacZ* transcriptional fusion were cultured in the presence (black bars) and presence (gray bars) of L-tartrate, and β-galactosidase activity quantified as described above. Bars represent geometric mean with geometric standard deviation. Each dot represents one biological replicate. WT; wild-type *, *P* < 0.05; *** *P* < 0.001; ns, not

### TtdV and TtdW bind to the *ttdU*-*ttdV* intergenic region

We next sought to uncover the mechanism TtdV and TtdW use to control the transcription of *ttdBAU*. We hypothesized that TtdV or TtdW could be binding to the region upstream of the *ttdU* gene to exert control. We cloned four biotinylated DNA probes that span the *ttdV*-*ttdU* intergenic region as well as part of the *ttdV* coding sequence (**Fig. 3A**) and purified C-terminal histidine-tagged TtdV (TtdV-His_6_) and TtdW (TtdW-His_6_) to assess protein-DNA interactions by electrophoretic mobility shift assays (EMSAs). Of the four probes tested, we observed a significant retardation when probe 4 was incubated with TtdW (**Fig. 3B**). We also detected more pronounced retardation of DNA probe 4 with increasing concentrations of TtdW (**Fig. 3C**). Competing labeled probe 4 with unlabeled probe 4 for TtdW binding resulted in the disappearance of the shift (**Fig. 3C**). TtdV did not bind to any of the probes (**Fig. 4A**). However, incubating TtdV, TtdW, and labeled probe 4 resulted in a new, slower migrating band, indicating that TtdV, TtdW and the DNA have formed a complex (**Fig. 4B**). Overall, the results from these experiments suggest that TtdW binds to a region of DNA closest to the *ttdU* gene and TtdV requires TtdW to form a complex with the target DNA.

**Figure 3:**
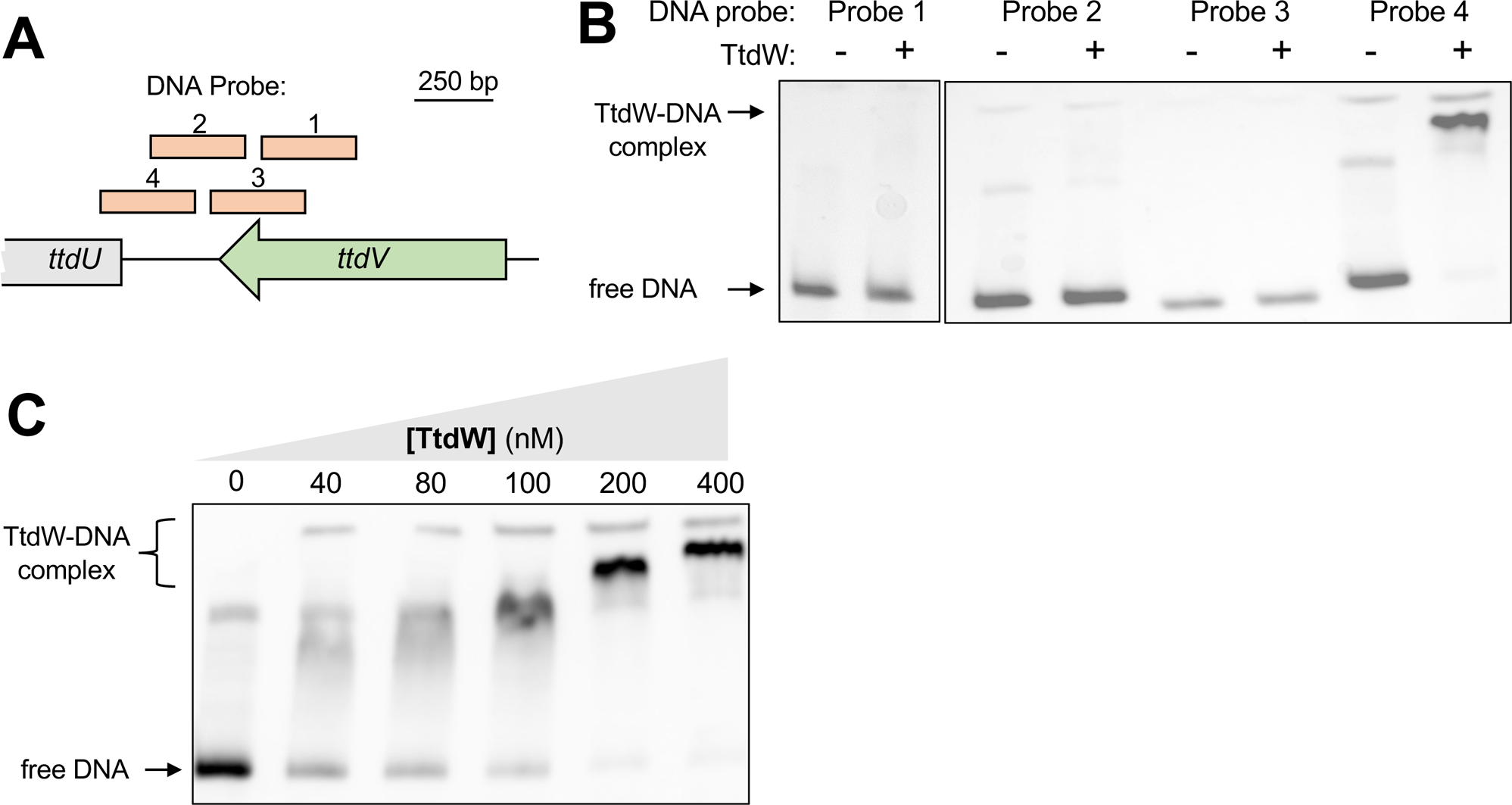
TtdW bind to the intergenic region between *ttdU* and *ttdV*. (**A**) Schematic representation of the DNA probes used to probe for interaction with TtdW. (**B – C**) Electrophoretic mobility shift assays contained 20 fmol of DNA probes and were resolved on a 6% 0.5X TBE polyacrylamide gel. TtdW-His_6_ was added at a concentration of 400 nM (**B**) or at increasing concentrations (**C**). Gel images are representative of 2 independent experiments. The approximate location of free, unbound DNA and a TtdW-DNA complex is indicated on the left side of the gel.

**Figure 4:**
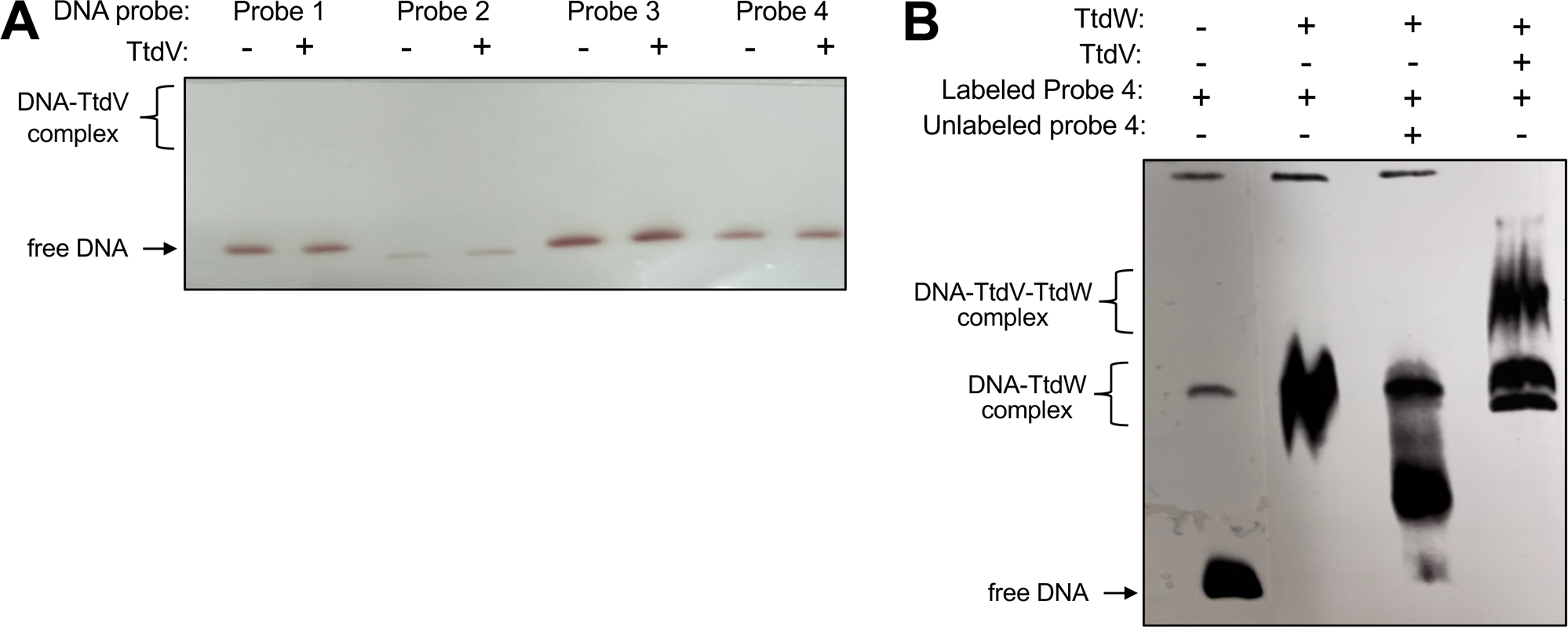
TtdV and TtdW bind to the *ttdU* - *ttdV* intergenic region. (**A**) Electrophoretic mobility shift assays contained 20 fmol of DNA probes and were resolved on a 6% 0.5x Tris-borate-EDTA polyacrylamide gel. TtdV-His_6_ was added at a concentration of 400 nM. (**B**) Electrophoretic mobility shift assays contained 20 fmol of DNA probes and were resolved on a 6% 0.5x Tris-borate-EDTA polyacrylamide gel. TtdV-His_6_ and TtdW-Hi_s6_ were added at a concentration of 400 nM as indicated. In one sample, a 100-fold excess of unlabeled probe 4 was added to the EMS binding reaction, as indicated. Gel images are representative of 2 independent experiments. The approximate location of free, unbound DNA and a protein-DNA complexes is indicated on the left side of the gel image.

### The ArcAB two-component system activates *ttdBAU* transcription

The ArcAB two-component system is a key regulatory system for fermentation and anaerobic respiration. ArcB, the sensor kinase, senses the redox state of the quinone pools that result from the cells metabolism, and phosphorylates ArcA, the response regulator, to control the expression of genes involved in energy metabolism of the cell (reviewed in (27)). In *E. coli*, ArcA was shown to have no effect on *ttdA* transcription in the absence of oxygen (28). However, it is plausible that ArcAB could influence the gene regulation for L-tartrate metabolism since the two-component system coordinates the directionality of the TCA cycle as the cell switches from fermentation to anaerobic respiration. During infection, the central metabolism switch of *Salmonella* switches as inflammation-derived electron acceptors become available during infection (reviewed in (19, 27)). We therefore hypothesized that the ArcAB two-component system is involved in regulating the L-tartrate utilization genes. We created strains that lack *arcA* or *arcB* and monitored *ttdA* transcription using a β-galactosidase reporter assay (**Fig. 5A**). The ArcAB system had no discernable effect on *ttdA* transcription in the absence of L-tartrate, however, induction of *ttdA* by L-tartrate was absent in both strains that lacked a functional ArcAB two-component system (**Fig. 5A**). Expression of recombinant ArcA from a plasmid in the *arcA* mutant restored responsiveness to L-tartrate (**Fig. S3**).

**Figure 5:**
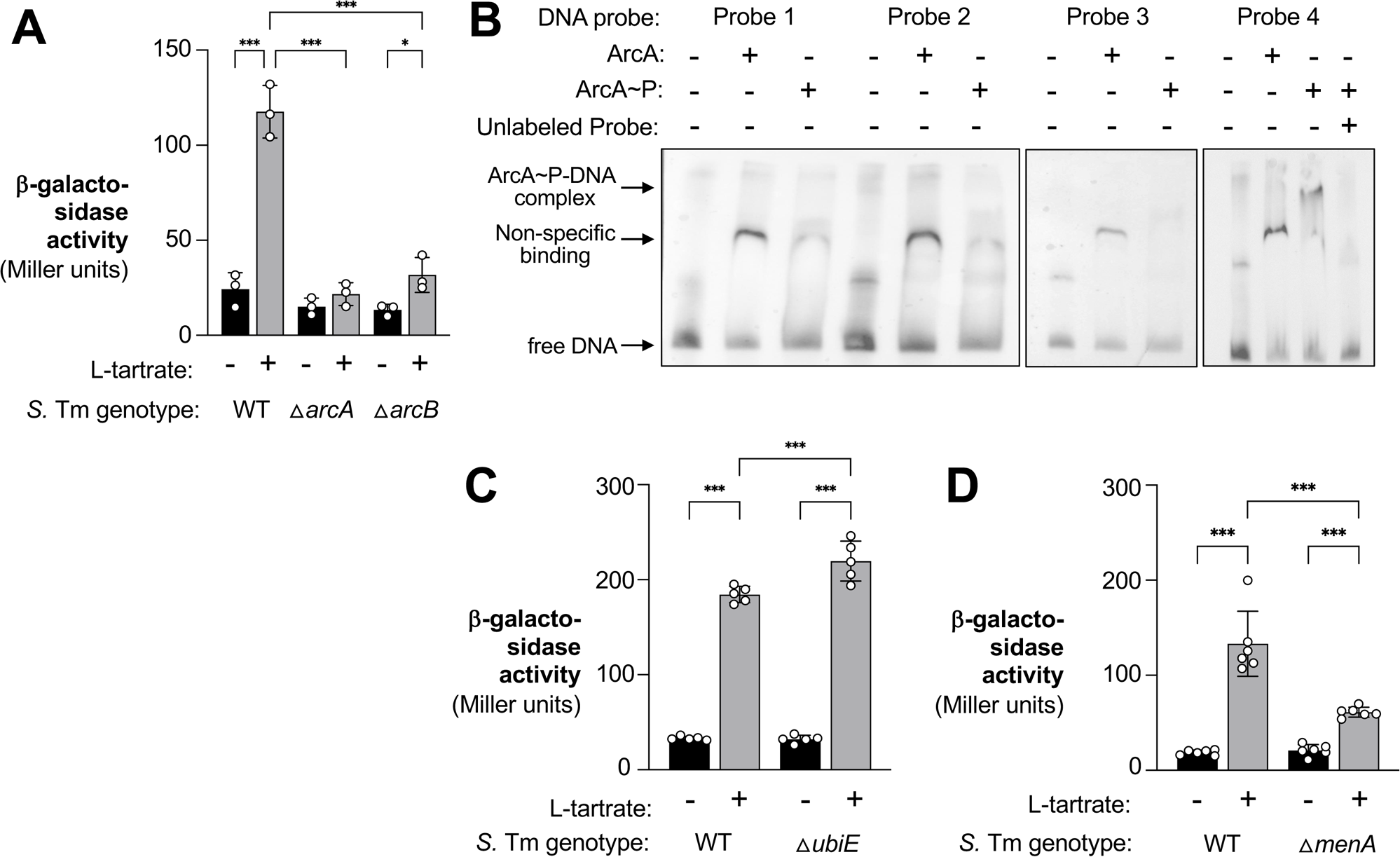
The ArcAB two-component system activates *ttdBAU* transcription. **(A)** *ttdA-lacZ* transcriptional reporter strains were anaerobically cultured in M9C media in the absence or presence of 0.1 mM L-tartrate. After grown for 3 hours, β-galactosidase activity was determined. The relevant genotype of each strain is shown below the graph. **(B)** Electrophoretic mobility shift (EMS) assays contained 50 fmol of DNA probes and were resolved on a 4.5% 0.5x Tris-borate-EDTA polyacrylamide gel. ArcA-His_6_ and phosphorylated ArcA-His_6_ were added at a concentration of 1500 nM, as indicated. A 50-fold excess amount of competitor unlabeled probe 4 to the EMS reaction mixture, as indicated. A 50-fold excess amount of competitor unlabeled probe 4 to the EMS reaction mixture, as indicated. The approximate location of free DNA and DNA-protein complexes is indicated on the left side of the gel image. **(C and D)** The *ttdA-lacZ* transcriptional reporter fusion strains, including a *ubiE* mutant **(C)** and a *menA* mutant **(D)**, were anaerobically cultured in M9C media in the absence or presence of 0.1 mM L-tartrate. After grown for 3 hours, β-galactosidase activity was determined. Bars represent geometric mean with geometric standard deviation. Each dot represents one biological replicate. WT; wild-type *, *P* < 0.05; *** *P* < 0.001; ns, not significant (ANOVA).

Next, we wanted to investigate if phosphorylated ArcA (ArcA∼P) would bind to the intergenic region between *ttdV* and *ttdU*. To this end, we purified a recombinant His_6_-ArcA fusion protein. We incubated the fusion protein with carbamoyl phosphate, a low molecular weight phospho-donor, before adding it to the EMSA binding reactions (**Fig. 5B**). Of note, all probes incubated with unphosphorylated ArcA exhibited a DNA shift, consistent with previous studies that have shown that ArcA binds non-specifically to DNA (29, 30). We found that ArcA∼P caused a specific retardation of probe 4. Addition of an excess of unlabeled probe 4 reversed the observed retardation of probe 4, indicating that the unlabeled probe 4 was competing with labeled probe 4 for ArcA∼P binding (**Fig. 5B**). These experiments suggest that, akin to TtdV and TtdW, ArcA∼P specifically binds to the *ttdV-ttdU* intergenic region.

During aerobic growth conditions, the ratio of ubiquinones (Q) to menaquinones (MK)/demethylmenaquinones (DMK) found in quinone pools are mostly comprised of Q and oxidize ArcB to silence its autophosphorylation activity, leading to ArcA inactivation. A shift from aerobic to anaerobic growth, however, triggers a change in the composition of the quinone pool in which MK/DMK make up a greater portion of the Q-MK/DMK ratio in the quinone pool. The predominance of MK/DMK activates ArcB kinase activity, leading to phosphorylation of ArcA (31, 32). We hypothesized that artificially changing the Q-MK/DMK ratio by inactivating Q or MK/DMK biosynthesis would lead to altered ArcB/ArcA regulation of the L-tartrate utilization genes. Inactivation of the *ubiE* gene results in the absence of Q and MK and an increased accumulation of DMK, and *menA* mutants cannot produce MK or DMK, but can synthesize Q (33). These strains were assessed for β-galactosidase activity under inducing and non-inducing conditions. In comparison to the L-tartrate induction of *ttdA* in the wild-type strain, the accumulation of DMK (*ubiE* mutant) resulted in enhanced transcription of *ttdA* in the presence of L-tartrate (**Fig. 5C**), consistent with the idea that a greater MK/DMK ratio will lead to activated ArcB sensing and ArcA activation and thus an increase in the transcription of *ttd* genes. In contrast, in a strain that lacks MK or DMK (*menA* mutant), the inducing effect L-tartrate had on *ttdA* transcription was drastically reduced (**Fig. 5D**), supporting the notion that an absence of these fermentation-associated quinones lead to silencing of the ArcAB two-component system and therefore cessation of transcription for the *ttd* operon. Our data suggest that activation of the L-tartrate gene cluster is dependent upon sensing of the redox state of the quinone pool.

### Regulation of L-tartrate metabolism is required for optimal fitness in the murine gut

During infection, L-tartrate is generated as a byproduct of reactive oxygen and reactive nitrogen metabolism (24). Since other electron acceptors, such as oxygen, nitrate, and tetrathionate become available during infection at later time points (16, 18, 34), we sought to clarify the link between these exogenous electron acceptors and L-tartrate utilization. Consistent with previous observations, anaerobic L-tartrate utilization ceases to provide a fitness advantage in the presence of alternative electron acceptors (**Fig. 6A**) (24). To determine whether these changes in L-tartrate utilization were driven by gene regulation, we cultured the *ttdA* transcriptional reporter strain in the presence of L-tartrate, and determined the impact of oxygen, nitrate, and tetrathionate on *ttdA* transcription (**Fig. 6B**). Consistent with our observation that electron acceptors interfere with the fitness advantage conferred by L-tartrate utilization, *ttdA* transcription was significantly repressed by all three exogenous electron acceptors.

**Figure 6:**
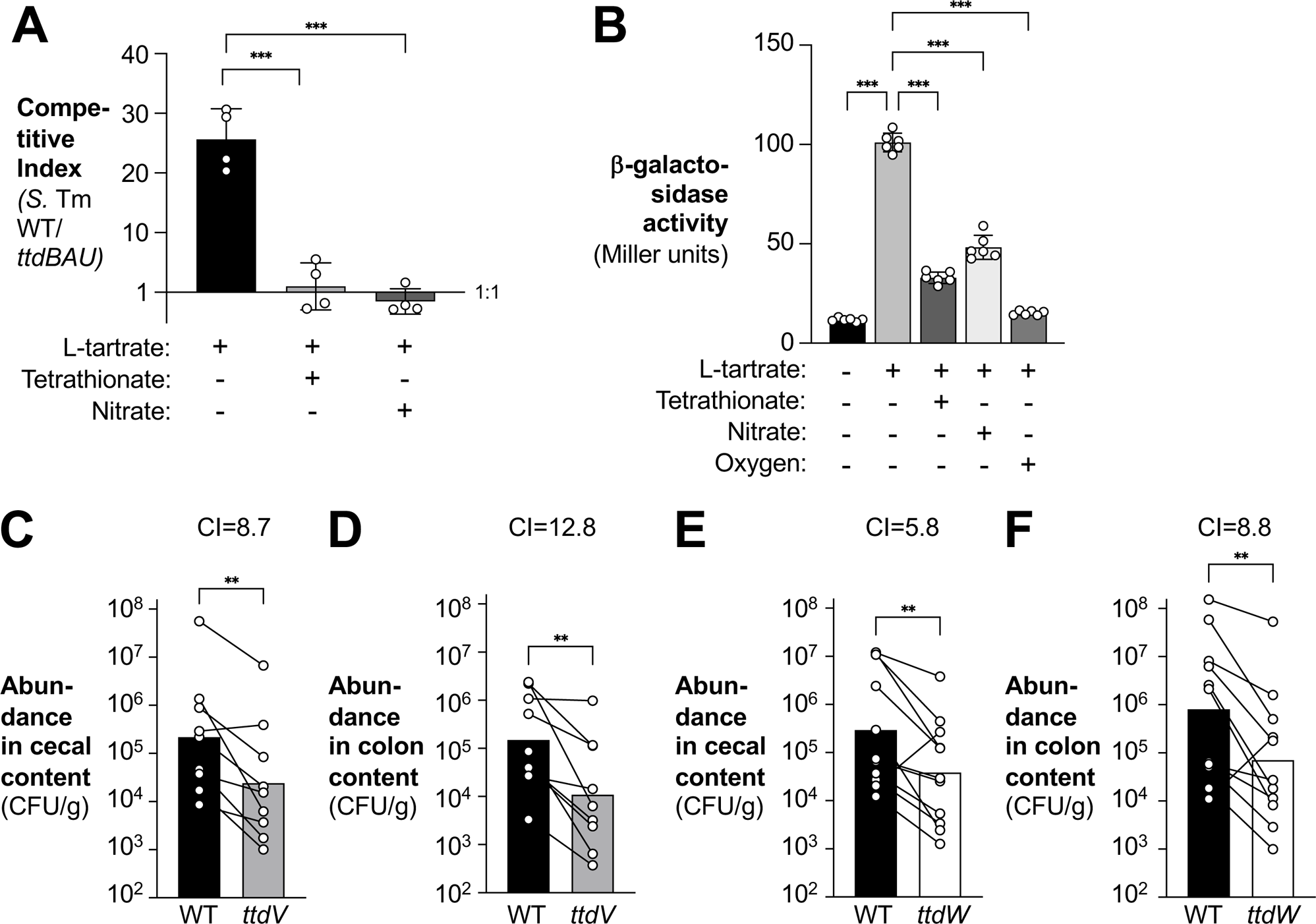
Regulation of L-tartrate metabolism is required for optimal fitness in the murine gut. **(A)** Minimal M9 media containing 20 mM L-tartrate, 20 mM glycerol, and 0.2 % casamino acids was supplemented with 40 mM tetrathionate or 40 mM nitrate as indicated. An equal mixture of the *S.* Tm wild-type strain (WT) and a Δ*ttdBAU* mutant strain were added and the suspension incubated in an anaerobic chamber for 18 h. The fitness of the strains in each condition was measured by determining the competitive index. **(B)** *ttdA-lacZ* transcriptional reporter strains were anaerobically cultured in M9C media in the absence or presence of 0.1 mM L-tartrate and 4 mM tetrathionate or 4 mM nitrate, and grown anaerobically or aerobically, as indicated. β-galactosidase activity was determined after 3 h of growth. (**C - F**) Groups of CBA mice were intragastrically inoculated with a 1:1 ratio of the *S.* Tm wild-type strain (black bars) and a *ttdV* mutant (gray bars)(**C** and **D**) or the wild-type strain (black bars) and a *ttdW* mutant (white bars)(**E** and **F**). Samples of the cecal (**C** and **E**) and colon content (**D** and **F**) were collected 4 days after infection and the fitness of each strain in the colon and cecal content was measured by determining the competitive index. Bars represent the geometric mean with geometric SD. Each dot represents one biological replicate. *, *P* < 0.05; **, *P* < 0.01; *** *P* < 0.001.

We next asked if the ability to regulate L-tartrate metabolism contributes to *Salmonella* fitness during infection. Groups of CBA mice were intragastrically inoculated with equal amounts of the *S*. Tm wild-type strain and a *ttdV* mutant, or the *S*. Tm wild-type strain and a *ttdW* mutant. After four days, colonic and cecal content were collected and plated on selective media (**Fig. 6C-F**). Fitness was assessed by calculating the ratio of the wild-type strain and the mutant in the contents, corrected by the corresponding ratio of the two strains in the inoculum (competitive index). Compared to the wild-type strain, the *ttdV* and *ttdW* mutants were less fit in the colon and cecum content, indicating that TtdV and TtdW-mediated gene regulation is required for optimal fitness of *S*. Tm.

## DISCUSSION

*S.* Tm has fine-tuned metabolism and virulence to cope with the transient niches it experiences during infection. Here, we investigated how L-tartrate, a metabolite generated as a byproduct of inflammatory ROS and RNS metabolism through the oxidative decarboxylation of monosaccharides, is sensed by *S*. Tm. Our *in vitro* studies enable us to formulate a specific model of how the L-tartrate utilization gene cluster is regulated (**Fig. 7**). Both TtdV and TtdW are negative regulators (**Fig. 2**), while the activated response regulator ArcA mediates positive regulation and is required for optimal transcription (**Fig. 5**). We propose that both TtdV and TtdW interact and that this complex represses transcription of the *ttdBAU* operon (**Fig. 3 and 4**). In the presence of L-tartrate, this repression is relieved (**Fig. 1**). Akin to TtdV and TtdW, ArcA∼P binds the *ttdV-ttdU* intergenic region (**Fig. 5**), and it is plausible that these two binding sites overlap. Our data are consistent with a scenario in which TtdV and TtdW prevent ArcA∼P from binding to the *ttdV-ttdU* intergenic region and prevent transcriptional activation by ArcA∼P. For substrate-induced activation, TtdV and TtdW are removed, possibly by L-tartrate binding to the complex, before ArcA∼P can activate transcription of *ttdBAU*. In the presence of respiratory electron acceptors, such as oxygen, unphosphorylated ArcA is unable to bind the *ttdV-ttdU* intergenic region, and transcription is minimal regardless of L-tartrate availability. One limitation of our study is that we were unable to conclusively demonstrate L-tartrate binding to purified TtdV and TtdW proteins. However, it is likely that L-tartrate, and not a downstream metabolite, is the actual cue to relieve TtdV and TtdW-mediated repression, since a mutant unable to degrade L-tartrate is fully responsive to L-tartrate-mediated induction.

**Figure 7:**
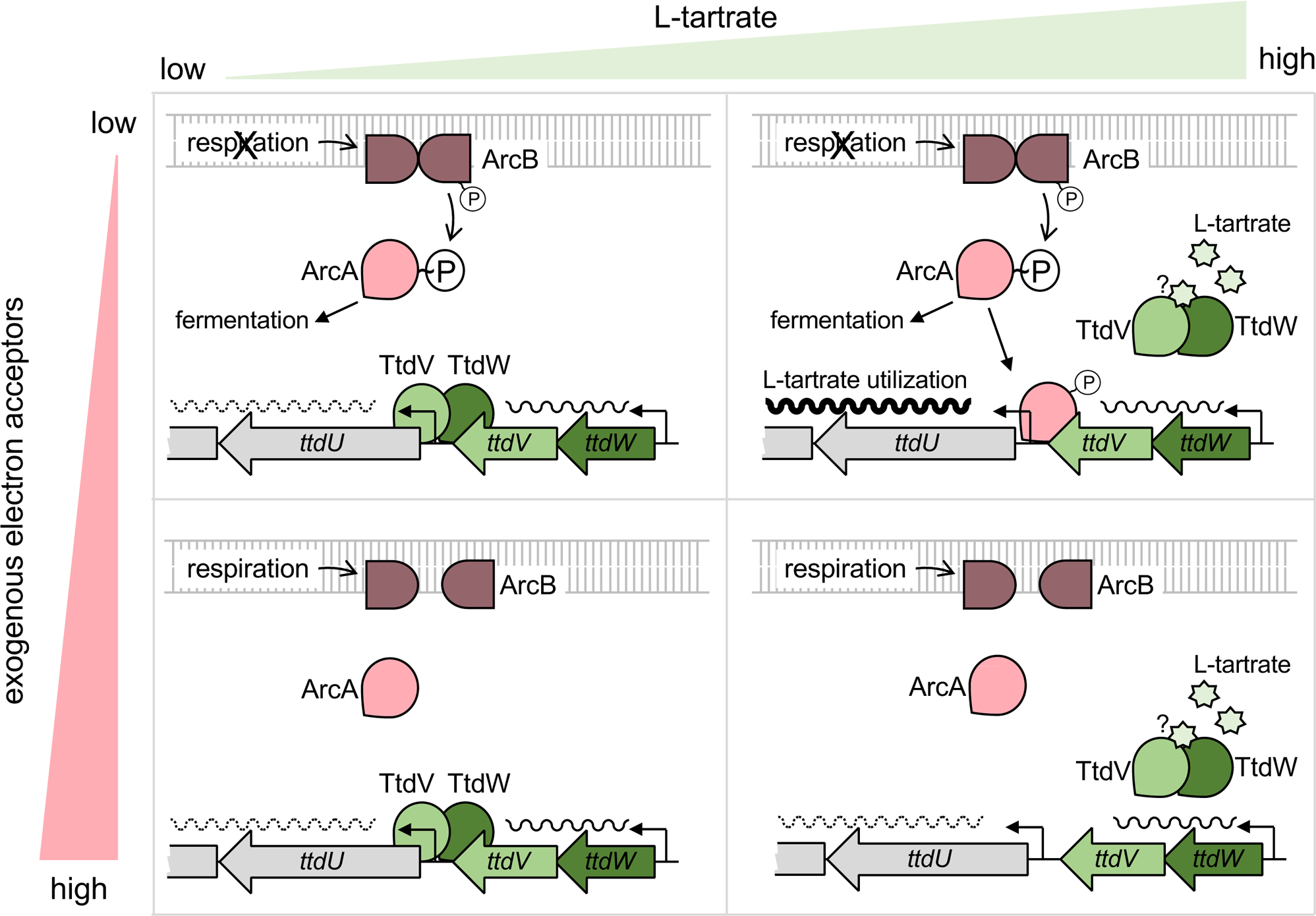
Proposed model for the regulation of the L-tartrate utilization gene cluster *ttdBAU ttdVW* in *S*. Tm. In the absence of exogenous electron acceptors and L-tartrate, TtdV and TtdW repress transcription of *ttdBAU* (top left panel). Increased availability of L-tartrate, in the absence of exogenous electron acceptors, leads to de-repression by TtdV and TtdW; phosphorylated ArcA binds in the ttdU-ttdV intergenic region and enhances transcription of *ttdBAU* (top right panel). In the presence of respiratory electron acceptors, the ArcAB system is inactive and unable to enhance transcription of *ttdBAU,* regardless of whether L-tartrate is absent (bottom left panel) or present (bottom right panel).

Our model (**Fig. 7**) illustrates how *Salmonella* integrates two infection-relevant cues, i.e. the availability of exogenous electron acceptors and the availability of the substrate, L-tartrate. We had previously shown that tartrate isomers, including L-tartrate, are generated as byproducts of inflammatory ROS and RNS metabolism, and that utilization of L-tartrate confers a fitness advantage early during infection (24). During that time, *S*. Tm utilizes L-tartrate early during infection in a branched TCA cycle, with a portion of L-tartrate being ultimately converted to fumarate to support fumarate respiration. At later time points, electron acceptors released by inflammatory macrophages and neutrophils, such as nitrate and tetrathionate, enable a population of *S*. Tm to respire and switch from a branched TCA cycle to a complete, oxidative TCA cycle (24). At this point, the fitness advantage conferred by L-tartrate utilization fades, likely due to repression of L-tartrate utilization genes by increased availability of electron acceptors (24) (**Fig. 6**). Only in the absence of energetically more favorable electron acceptors, and when the substrate is present at appreciable concentrations, genes involved in L-tartrate utilization are optimally transcribed. These data could explain why L-tartrate is primarily converted to fumarate for fumarate reduction, and L-tartrate conversion to oxaloacetate is not used as an anaplerotic reaction. In the presence of exogenous electron acceptors and more efficient electron transport chains, fumarate reduction becomes superfluous and L-tartrate degradation via the *ttdBAU* gene cluster ceases due to gene repression.

Regulation of gene expression is critical for *Salmonella* to establish infection. Many of the cues involved in virulence gene regulation are metabolites as well as host-derived antimicrobials. Metabolites regulate the expression of T3SS machinery to minimize any metabolic cost of expressing while enabling tissue-specific invasion and tissue replication (35, 36). Our data suggest that TtdV and TtdW, two regulatory proteins required for repression of genes involved in L-tartrate utilization, are required for optimal fitness during gut colonization. Regardless of whether the cell exhibits a branched or complete TCA cycle, the central metabolism fulfills key physiological functions, i.e. redox balance, interconversion of metabolic building blocks, and energy generation. As such, the TCA cycle, as well as major pathways connecting to it, are subject to complex control mechanisms (37). TtdAB converts L-tartrate directly to the TCA cycle intermediate oxaloacetate, thus likely requiring tight regulation of TtdAB activity. In the absence of TtdV- and TtdW-mediated repression, uncontrolled uptake of L-tartrate and immediate conversion to oxaloacetate could perturb metabolic flux in the TCA cycle, thus leading to slower growth and decreased gut colonization (**Fig. 6**).

L-tartrate metabolism has been studied in detail in commensal *E. coli* (28, 38, 39). The L-tartrate utilization gene cluster in *E. coli* encodes a single LysR-type transcriptional regulator TtdR, whereas the presence of a second transcriptional regulator in *S*. Tm deviates from the genetic structure of the operon found in *E*. *coli*. The primary amino acid sequences of TtdR (*E. coli*), TtdV (*S.* Tm), and TtdW (*S.* Tm) exhibit no apparent similarities, suggesting that gene regulation of L-tartrate utilization differs between *E. coli* and *Salmonella*. Furthermore, ArcAB regulates L-tartrate utilization in *S*. Tm (**Fig. 5**), but not commensal *E. coli* (28). These observations raise the possibility that the L-tartrate gene regulation in *Salmonella* could be an adaptation to the pathogenic lifecycle of *Salmonella*.

## MATERIALS AND METHODS

### Bacterial culture

Unless indicated otherwise, *S*. Tm and *E. coli* strains were grown on LB plates (10 g/l tryptone, 5 g/l yeast extract, 10 g/l sodium chloride, 15 g/l agar) and LB broth (10 g/l tryptone, 5 g/l yeast extract, 10 g/l sodium chloride) under aerobic conditions at 37°C for 16 hours. Agar plates were supplemented with kanamycin (Kan), nalidixic acid (Nal), carbenicillin (Carb), or chloramphenicol (Cm) at concentrations 50 μg/ml, 50 μg/ml, 100 μg/ml, and 15 μg/ml, respectively. The chromogenic molecule X-Phos (5-bromo-4-chloro-3-indolyl phosphate, Chem-Impex) was added to agar plates with the purpose of detecting *S*. Tm strains that express the acidic phosphatase PhoN. The base media used in all *in vitro* studies contained M9 minimal salts (0.068 g/l Na_2_HPO_4_*7H_2_O, 0.03 g/l KH_2_PO_4_, 0.005 g/l NaCl, 0.01 g/l NH_4_Cl), 18.5 mM Bacto casamino acids and 85.5 mM sodium chloride.

### Plasmids, bacterial strains, and generation of mutants

The bacterial strains and plasmids used in this study are found in Table 1 and all primers listed in Table 2. The upstream and downstream regions of *ttdV*, *ttdW*, *ubiE,* and *menA* were amplified by PCR from the IR715 genome using Q5 High Fidelity Master Mix (New England Biolabs). PCR products were purified using the QIAEX II Gel Extraction Kit (Qiagen). Using Gibson Assembly (New England Biolabs), the upstream and downstream regions of *ttdV*, *ttdW*, *ttdVW,* and *ttBAUVW* were inserted into SphI-digested pGP706 and the upstream and downstream regions of *ubiE* and *menA* were inserted into SphI-digested pRDH10, giving rise to pSC1, pSC2, pSC3, pVR2, pMW31, and pMW35, respectively. *E. coli* DH5α λ*pir* served as the cloning strain for all suicide plasmids. The DNA sequence of upstream and downstream regions cloned into the suicide plasmid pGP706 was verified by Sanger sequencing. Suicide plasmids were introduced into *S*. Tm via conjugation. *E. coli* S17-1 λ*pir* served as the donor strain for conjugation. Exconjugants (merodiploid) were selected on LB agar plates supplemented with Nal and Kan. To perform counterselection, a merodiploid colony was grown overnight in LB broth at 37°C with aeration and a diluted samples of this culture plated on sucrose plates (5% sucrose, 15 g/l agar, 8 g/l nutrient broth base). Clean deletions were verified via PCR using Quick-Load Taq Master Mix (New England Biolabs) or Q5 High Fidelity Master Mix. This cloning strategy was used to make bacterial strains SC1 (Δ*ttdV*), SC2 (Δ*ttdW*), SC3 (Δ*ttdVW*), MW201 (Δ*menA*), MW208 (Δ*ubiE*), and VR2 (Δ*ttdBAUVW*).

To create complementation strains VR3, VR29 and VR28, we first created pPhoKm. We designed pPhoKm by inserting PCR amplified upstream and downstream DNA regions of the neutral *phoN* locus that would remove the existing SphI cut site from the plasmid and introduce a new SphI cut site in between the two *phoN* flanking regions and inserted these modified flanking regions into SphI-digested pGP706 using Gibson Assembly. We then PCR amplified the native promoter and gene(s) of interest and inserted the fragment into SphI-digested pPhoKm. We used the same strategy of making clean deletions to make clean insertions of the promoter and gene of interest in the *phoN* region.

To generate lacZ reporter strains, we constructed pVR7. To this end, 330 bp from the 3’ end of the *ttdA* gene were PCR amplified, purified, and cloned into SmaI-digested pFUSE. S17-1 λ*pir* harboring pVR7 was conjugated with *S.* Tm strain IR715 and exconjugants were selected for on LB plates supplemented with Nal and Cm. This cloning strategy created bacterial strain VR4 (*ttdA*::pVR7). Conjugation of pVR7 into SC1, SC2, SC3, VR3, VR28, VR29 resulted in transcriptional fusion strains VR9, VR10, VR11, VR35, VR36, and VR34, respectively. The *ttdA*::pVR7 mutation was transduced by phage P22 (40) into MW118 (Δ*arcB*), MW119 (Δ*arcA*), MW201, MW208, to generate VR15, VR14, VR50, VR48, respectively.

### β-Galactosidase Assay

Media prepared for all β-Galactosidase assays that included L-tartrate were supplemented with a final concentration of 0.1 mM L-tartrate, unless indicated otherwise. Potassium tetrathionate (Sigma) or sodium nitrate (Sigma) were added at a final concentration of 4 mM. Three milliliters of media were preincubated in the anaerobic chamber (5% hydrogen, 5% CO_2_, 90% nitrogen) 1 d prior to inoculation. Overnight cultures grown aerobically in LB broth and Cm were sub-cultured in the pre-reduced media at a 1:20 dilution and incubated in the anaerobic chamber at 37°C for 3 h. The sub-cultures were taken out of the anaerobic chamber and placed on ice for 20 min. The protocol described by Miller (41) was used to perform the assays. Briefly, after the optical density at a wavelength of 600 nm (OD_600_) was measured, 100 μl or 500 μl of the sub-culture was combined with 900 μl or 500 μl sterile PBS supplemented with β-mercaptoethanol (Thermo Fisher), respectively. Bacterial lysis was carried out by adding 40 μl chloroform and 20 μl 0.1% SDS, then vortexed for 10 s, followed by a 5 min incubation at room temperature. To begin the reaction, 200 μl of 4 mg/ml ONPG (ortho-nitrophenyl-β-galactoside, Thermo Scientific) was added, and after allowing the reaction sufficient time to develop a yellow color, the reaction was stopped by adding 500 μl of 1 M Na_2_CO_3_ and a final mix (vortex). The absorbance at a wavelength of 420 nm and 550 nm was measured and the Miller units were calculated as follows: Miller Units = 1,000 x ([A_420_ – (1.75 x A_550_)]/(t x v x OD_600_)), where t is the time in minutes and v is the volume is milliliters.

### Quantification of mRNA in *S.* Tm grown *in vitro*

Nine milliliters of freshly prepared media were placed in the anaerobic chamber 1 d prior to inoculation. Overnight cultures grown with aeration in LB broth were sub-cultured in the pre-reduced media at a 1:10 dilution then incubated at 37°C for 3 h. At the end of the incubation, the cultures were removed from the anaerobic chamber, placed in a 4°C centrifuge and the suspension centrifuged for 5 min at 3220 g. After discarding the media, total RNA was extracted using the Aurum Total RNA Mini Kit (Bio-Rad), treated with DNase I (Thermo Fisher) twice then stored at −80°C.

### RT-PCR and qPCR

The concentration of the DNase treated RNA was determined by measuring the A_260_/A_280_ ratio (Nanodrop One, Thermo Scientific). A 25 μl reaction was prepared for each sample using the following TaqMan reverse transcription reagents: 2.5 μl 10X RT PCR buffer, 5.5 μl 25 mM MgCl_2_, 5 μl dNTP mix, 2.5 mM each, 1.3 μl of 50 μM random hexamers, 0.5 μl RNase inhibitor, and 0.6 μl reverse transcriptase (RT) and 9.6 μl of template RNA. To account for DNA contamination, control reactions lacking RT were prepared for each sample by mixing RT-PCR buffer, template RNA and water. Samples were incubated in a thermal cycler at 25 °C for 10 min, 48 °C for 30 min, 95 °C for 5 min.

To perform real-time PCR (qPCR), we prepared a 25 μl reaction for each sample: 12.5 μl SYBR Green Mix, 0.625 μl of forward and reverse primer, 7.25 μl water and 4 μl cDNA. Primer sequences are listed in Table 2. Two technical replicates for each biological replicate were assayed using the SYBR green qPCR supplied protocol from QuantStudio 6 Real-Time System. QuantStudio qPCR software was used for data analysis, determining baselines through the baseline threshold algorithm. Subsequently, Microsoft Excel and Prism 10 was used to further analyze the data, employing the comparative threshold cycle method.

### Protein expression and purification

The strains and plasmids used to generate and maintain the recombinant proteins are described in Table 1 and Table 2, respectively. To generate pVR17 and pVR18, we PCR amplified the *ttdV* and *ttdW* genes, leaving out the stop codon, from genomic material of *S*. Tm IR715 as the template and using Q5 High Fidelity Master Mix. The PCR products were purified using the same DNA purification kit for generating mutant strains, and Gibson assembled with XhoI-digested pET21(+). The Gibson Assembly reaction was transformed into *E. coli* DH5α λ*pir*, and the transformed bacteria were plated on LB agar plates supplemented with Carb. The plasmids were confirmed via PCR and subsequently cultured in 100 ml LB broth with 100 μl Carb overnight to extract the recombinant vectors using the Qiagen Plasmid Midi Kit. pVR17 and pVR18 were used to transform *E. coli* BL21 (DE3) cells and the transformants were plated on LB agar with Carb. The presence of the recombinant plasmids was confirmed by PCR.

To express the recombinant proteins from BL21 cells, we inoculated 10 ml LB broth supplemented with 1 M glucose for a final concentration of 20 mM and 10 μl Carb with one colony of the BL21 strains harboring the recombinant plasmids pVR17 and pVR18. The starter culture was grown overnight at 37°C with aeration. The following day, 5 ml of the starter cultures were aliquoted into 100 ml LB broth with 100 μl Carb. The sub-culture was grown at 37°C with vigorous shaking for about 1 h or until the OD_600_ range was between 0.5 and 0.7. The sub-culture was then supplemented with 100 μl of a 1M ITPG stock for a final concentration of 1 mM to induce protein expression of recombinant TtdV and TtdW. Protein induction lasted for 4-5 h at 37°C shaking at 250 rpm. The sub-culture was split into 50 ml tubes and centrifuged at 4°C at 3220 g for 20 min. The supernatant was discarded, and the cell pellets were stored at −20°C until ready for further extraction. The cell pellets were thawed on ice for 15 min before adding 5 ml of ice-cold lysis buffer (50 mM NaH_2_PO_4_, 300 mM NaCl, 10 mM imidazole) and 5 mg of lysozyme for a final concentration of 1 mg/ml was added to the lysate and incubated on ice for 30 min. To lyse the cells, the 5 ml cell suspension was kept on ice and sonicated with a microtip with 10 s bursts and 10 s rest periods for 6 times. The cleared lysate was centrifuged at 10,000 g for 20 min at 4°C. The supernatant was decanted into a new 15 ml tube (soluble fraction) and the pellet was resuspended with 5 ml lysis buffer (insoluble fraction). The whole cell lysates pre- and post IPTG induction and cellular fractions were loaded on a 10 % polyacrylamide gel and Coomassie stained to assess expression and solubility of recombinant TtdV-His_6_ or TtdW-His_6_. Recombinant TtdV-His_6_ or TtdW-His_6_ were both found in the soluble fraction when expressed from pET21(+) vector. To isolate recombinant protein from the soluble fraction, we followed the vendor supplied protocol of the His-Pur Ni-NTA Purification Kit (3 ml, Thermo Fisher). The flow through, wash and elution fractions were analyzed on a 10% polyacrylamide gel that was Coomassie stained. We used the Pierce BCA Protein Assay Kit (Thermo Fisher) to measure the protein concentration from all three elution fractions. The third elution fraction had the least amount of protein contamination and was stored at 4°C and used for subsequent EMSA reactions.

To generate pVR55, we restriction digested pET14b with BamHI, resolved the digested vector on a 0.7% agarose gel and purified the DNA using the QIAEX II Gel Extraction Kit. We PCR amplified the *arcA* gene, leaving out the start codon, from the *S*. Tm IR715 genome. After agarose gel electrophoresis, DNA was extracted and Gibson Assembly performed with BamHI-digested pET14b as described above. Next, we transformed *E. coli* DH5α λ*pir* with 2 μl of the Gibson Assembly and plated the transformants on LB agar supplemented with Carb. We verified the insertion of *arcA* into pET14b by PCR, then grew a single colony in 100 m LB broth supplemented with 100 μl Carb overnight to prepare for plasmid extraction. We used the Qiagen Plasmid Midi Kit to extract the plasmid. We used *E. coli* BL21 pLysE strain for expression of *arcA* from pET14b due to potential toxic side effects that recombinant ArcA would have on the endogenous ArcAB sensing. We plated the electroporated cells onto LB agar with Carb. Colonies were purified and confirmed to harbor the plasmid via PCR.

To purify His_6_-ArcA, we grew a 6 ml starter culture with LB broth, 6 μl Carb, 3 μl Cm and 120 μl 1 M glucose (final concentration 20 mM) overnight at 37°C with aeration. We then sub-cultured 5 ml of the overnight starter culture into 250 ml LB broth containing 250 μl Carb. The sub-culture was shaken at 250 rpm at 37°C until the culture reached an optical density (OD_600_) of 0.4-0.6. The sub-culture was induced with 250 μl of 1 M ITPG (final concentration 1 mM) and kept shaking for 5 h. One milliliter aliquots were taken before and after induction to serve as whole cell lysates controls. The sub-culture was split into two 250 ml centrifuge bottles then centrifuged at 3220 g for 20 min at 4°C. The supernatant was removed, and pellets stored at −20°C until the next day. The pellets were thawed on ice for 15 min before resuspending with 7 ml of ice-cold lysis buffer, collecting the pellets into one 15 ml tubes then adding 7 mg of lysozyme. The pellets were kept on ice for 30 min then sonicated with a microtip 6 times with 10 s bursts and 10 s rests. The cleared lysate was centrifuged at 10,000 g for 20 min at 4°C, the supernatant (soluble fraction) was decanted into a new 15 ml tube and the pellet (insoluble fraction) was resuspended in 7 ml lysis buffer. The whole cell lysates from the uninduced and induced culture and the fractions were analyzed via SDS-PAGE and Coomassie stained to verify His_6_-ArcA expression and that it was found in the soluble fraction. We used the same protocol and kits to purify His_6_-ArcA and measure each fractions’ concentration.

### Electrophoretic Mobility Shift Assay (EMSA) using TtdV and TtdW

We prepared the biotinylated (labeled) and unbiotinylated (unlabeled) DNA probes by PCR amplification using *S*. Tm IR715 genomic DNA as the template and Q5 High Fidelity Master Mix using the following thermocycling profile: denature at 98°C for 1 min, 35 cycles of melting at 98°C for 10 s, annealing at 50-72°C for 30 s, and extension at 72°C for 20 s with a final extension of 72°C for 1 min and an infinite hold at 4°C. The PCR products were loaded onto a 1% agarose gel, excised, and purified using the QIAEX II Gel Extraction Kit. We used the LightShift Chemiluminescent EMSA Kit (Thermo Scientific) and protocol for all EMSA reactions. For the competition reactions, we incubated unlabeled probe 4 with TtdW-His_6_ for 5 min at room temperature before adding labeled probe 4. After a 20 min incubation at room temperature, we loaded the EMSA reactions onto TBE gels (0.5x TBE buffer, 4.5 – 6% polyacrylamide, 0.5% glycerol, 0.06% ammonium persulfate, 0.2% tetramethylethylenediamine). TBE buffer contained 5.4 g/l Tris base, 2.8 g/l boric acid, and 10 mM EDTA. To transfer the binding reactions to the nylon membrane (Biodyne™ B Nylon Membrane, Thermo Scientific), we semi-dry transferred the reactions at 15 V for 45 min and immediately crosslinked the transferred DNA to the membrane using a commercial UV-light crosslinking oven (UV Stratalinker, Stratagene) with a 45-60 s exposure using the auto crosslink function.

### *In vitro* ArcA phosphorylation reaction and DNA binding assays using ArcA and ArcA∼P

To phosphorylate ArcA before adding the modified protein to the EMSA reactions, we incubated purified His_6_-ArcA at a final concentration of 80 μg/ml, with carbamoyl phosphate (final concentration 50 mM) and phosphorylation buffer (50 mM Tris-Cl, pH 7.0, 50 mM KCl, 20 mM MgCl_2_, 20 mM dithiothreitol, 10 μg/m bovine serum albumin, all final concentrations) for a 60 μl reaction. After a 1 h incubation at room temperature, the phosphorylation reaction was stopped with 200 mM EDTA pH 8.0. The phosphorylated ArcA was then added to the appropriate EMSA reactions.

The same probes were used to assess ArcA/ArcA∼P binding, and the same kit and protocol were used to perform all EMSA reactions, with minor modifications. We included 10 mM EDTA to the binding reactions. To assess competing DNA binding between labeled probe 4 and unlabeled probe 4, we incubated ArcA∼P with unlabeled probe 4 for 5 min at room temperature before adding labeled probe 4. The EMSA reactions were loaded onto a 4.5% 0.5X TBE gel and the same transfer and crosslinking process was followed.

### Anaerobic growth of *S.* Tm in M9 media supplemented with casamino acids

Media that was pre incubated in the anaerobic chamber for 1 d was inoculated with equal ratios of AJB715 and LS25 at a final concentration of 1 × 10^3^ CFU/ml of each strain. Final concentrations of 20 mM L-tartrate, 20 mM glycerol, and 0.2% casamino acids, as well as 40 mM sodium nitrate or 40 mM potassium tetrathionate were added as indicated. The strains were grown in the anaerobic chamber for 18 h at 37°C and recovery of each strain were determined by spread plating serial dilutions on selective LB agar plates.

### CBA infection model

All mouse experiments were performed in accordance with the Institutional Animal Care and Use Committees at UT Southwestern Medical Center and at UC Davis. Groups of CBA mice (aged 8-10 weeks old) were intragastrically infected with 5 × 10^8^ CFU of each strain for competitive infection experiments. After 4 days, colonic and cecal luminal content were collected for bacterial enumeration. Serial dilutions of intestinal content were plated on selective media.

## Funding

Work in S.E.W.’s lab was funded by the NIH (AI118807, AI66263, AI171537), the Burroughs Wellcome Fund (1017880), and a grant from the US - Israel Binational Science Foundation (2021025). Any opinions, findings, and conclusions or recommendations expressed in this material are those of the author(s) and do not necessarily reflect the views of the funding agencies. The funders had no role in study design, data collection and interpretation, or the decision to submit the work for publication.

## Author contributions

VKR, MGW, AGJ, NWT, and LS. performed and analyzed the *in vivo* and *in vitro* experiments. SC contributed critical tools. VKR, AGJ, DRH, and SEW designed the experiments, interpreted the data, and wrote the manuscript with contributions from all authors.

## Data and materials availability

Materials transfer agreements and regulatory permits may be required to transfer materials described in this study.

